# Phycocyanobilin biosynthesis in *Galdieria sulphuraria* requires isomerization of phycoerythrobilin synthesized by bilin reductases

**DOI:** 10.1101/2025.04.02.646756

**Authors:** Federica Frascogna, Nathan C. Rockwell, Jana Hartmann, Julie Mudler, Nicole Frankenberg-Dinkel

**Affiliations:** Department of Microbiology, RPTU Kaiserslautern-Landau, 67663 Kaiserslautern, Germany; Department of Molecular and Cellular Biology, One Shields Avenue, University of California at Davis, Davis, CA 95616 USA

## Abstract

Cyanobacteria and red algae (Rhodophyta) utilize phycobilisomes as light-harvesting antennae to carry out oxygenic photosynthesis. Phycobilisomes are large protein complexes comprised of phycobiliproteins and associated linker proteins, with phycobiliproteins carrying one or more linear tetrapyrrole (bilin) chromophores. Bilin biosynthesis is typically catalyzed by ferredoxin-dependent bilin reductases (FDBRs). Cyanobacterial genomes encode one FDBR, PcyA, for biosynthesis of phycocyanobilin (PCB), which is required both for harvesting red light and for coupling phycobilisomes to photosynthetic reaction centers via energy transfer. Some cyanobacteria also contain two additional FDBRs (PebA and PebB) for synthesis of the green-absorbing phycoerythrobilin (PEB). Most rhodophytes contain only PebA and PebB homologs (PEBA and PEBB), so the biosynthesis of PCB in such organisms is not understood. This process is especially relevant in the early-branching rhodophyte Galdieria sulphuraria, which contains PCB as the main light-harvesting chromophore yet contains the PEBA and PEBB FDBRs for PEB biosynthesis. In this organism, PCB biosynthesis is thought to be dependent both on FDBRs and on a yet-to-be identified bilin isomerase. Our current work reaffirms these findings. Phylogenetic analysis clearly places G. sulphuraria FDBRs in the PEBA and PEBB clades, in contrast to the PCYA found in Cyanidioschyzon merolae, another early-branching rhodophyte. All three rhodophyte FDBRs were found to have the expected substrate preferences and enzymatic activities after recombinant expression, affinity purification, and in vitro characterization. Therefore, bilin biosynthesis in G. sulphuraria results in biosynthesis of PEB rather than PCB. However, a previous procedure allowed partial purification of a PEB:PCB isomerase, clearly demonstrating the existence of such an activity and adding to our understanding of bilin biosynthesis and algal light harvesting.

## Introduction

Phycobiliproteins (PBPs) are a family of fluorescent proteins primarily found in cyanobacteria, red algae, glaucophytes and cryptophytes, where they play a crucial role in light-harvesting during photosynthesis (Glazer, 1994; Croce & Van Amerongen, 2014; MacColl &Guard-Friar, 2018; Li *et al*., 2019). PBPs enhance the light-harvesting machinery of these organisms by absorbing light in the 500-650 nm range, the so called “Green gap” where chlorophyll poorly absorbs (Green & Parson, 2003; Blankenship, 2008; Polívka & Frank, 2010; Chen & Blankenship, 2011). These proteins are most often assembled into large complexes known as phycobilisomes (PBSs), which transfer the captured light energy to both photosystem II and photosystem I reaction centers (Gantt & Conti, 1966; Glazer, 1977; Grossman *et al*., 1993; Watanabe & Ikeuchi, 2013). The ability of PBPs to harvest light is dependent on open-chain tetrapyrrole chromophores, called phycobilins (or short bilins). For a long time, the early-branching Rhodophytes of the Cyanidiales served as model organisms to study the biosynthesis of these chromophores. The first studies on *Galdieria sulphuraria* (formerly mistaken for *Cyanidium caldarium*) (De Luca & Taddei, 1976; Merola *et al*., 1981; Gross & Schnarrenberger, 1995; Albertano *et al*., 2000; Cozzolino *et al*., 2000) implied that the open-chain tetrapyrrole biliverdin IXα (BV) is the precursor of all phycobilins (Beale & Cornejo, 1983, 1984). In subsequent studies, BV was shown to be reduced by an unknown ferredoxin-dependent enzyme to phycoerythrobilin (PEB), and then converted, via isomerization, to phycocyanobilin (PCB) (Beale & Cornejo, 1991a, b, c). Around a decade later, the enzymes responsible for BV reduction were identified as the so-called ferredoxin-dependent bilin reductases (FDBRs) (Frankenberg *et al*., 2001; McDowell & Lagarias, 2001). In general, the first universal step in the biosynthesis of bilins is the ferredoxin-dependent oxidation of heme at the α-methine bridge by heme oxygenase (HO) (Ortiz De Montellano, 2000; Yoshida & Migita, 2000; Wilks, 2002). This reaction yields BV and additionally liberates H_2_O, CO, and Fe^2+^ (Cornejo & Beale, 1988; Kikuchi *et al*., 2005). BV is subsequently reduced to specific bilins by FDBRs (Supporting Figure S1) (Frankenberg *et al*., 2001; Frankenberg & Lagarias, 2003a, b; Dammeyer & Frankenberg-Dinkel, 2008; Ledermann *et al*., 2018). Although Beale & Cornejo showed that PCB production occurs via PEB in *G. sulphuraria*, an enzyme directly reducing BV to PCB, named phycocyanobilin oxidoreductase (PcyA), was subsequently discovered in cyanobacteria (Frankenberg *et al*., 2001; Frankenberg & Lagarias, 2003a). In cyanobacteria, some other bacteria, glaucophyte algae, and a small subset of cryptophyte algae, PcyA is responsible for the 4-electron reduction of BV to PCB via 18^1^,18^2^-dihydrobiliverdin (18^1^,18^2^-DHBV) (Frankenberg & Lagarias, 2003b; Tu *et al*., 2004; Hagiwara *et al*., 2006; Hagiwara *et al*., 2010; Tu *et al*., 2007). More recently, PCB was found to be synthesized in a similar fashion by the FDBR HY2 of streptophyte algae, with the possible occurrence of either 18^1^,18^2^-DHBV or phytochromobilin (PФB) as the biosynthetic intermediate (Rockwell *et al*., 2017; Frascogna *et al*., 2023). However, in these organisms, PCB is bound to phytochromes and used for light sensing rather than light harvesting (Rockwell *et al*., 2017). Based on spectroscopic data, the PBSs of *Galdieria* spp., as in other Cyanidiales, are only believed to be composed of an allophycocyanin (APC) core and peripheral rods containing phycocyanin (PC), both binding PCB (Moon *et al*., 2014; Sarian *et al*., 2016; Rahman *et al*., 2017; Ferraro *et al*., 2020; Sommer *et al*., 2021). Therefore, the occurrence of PEB biosynthesis in the absence of detectable PEB chromophore and the requirement for PCB chromophore in the absence of PCB biosynthesis presents an interesting conundrum.

In this study, a combination of *in silico* approaches and enzymology was used to investigate PCB biosynthesis in *G. sulphuraria*. First, phylogenetic analysis reaffirmed the placement of FDBRs from *G. sulphuraria* and its relative *Cyanidioschyzon merolae* in the expected clades while providing further insight into the distribution of FDBR lineages in different rhodophytes and cryptophytes. These enzymes were then proven to function according to their phylogenetic classification after recombinant expression. In particular, *G. sulphuraria* was shown to possess only the FDBRs involved in PEB production, namely PEBA (*Gs*PEBA) and PEBB (*Gs*PEBB), reinforcing the isomerase hypothesis of Beale & Cornejo, while *C. merolae* is instead endowed with PCYA only (*Cm*PCYA). Finally, an *in vivo* approach, based on the method of Beale & Cornejo, was used to enrich an isomerase-active protein fraction of *G. sulphuraria* (Beale & Cornejo, 1991b). However, efforts to identify this putative enzyme via mass spectrometry proved to be intractable due to the high protein diversity of the sample and the overall poor proteomic annotation, requiring further investigation in the future. This unusual pathway may also provide insights into the broader evolutionary trajectory of photosynthetic pigment synthesis that stemmed from the primary endosymbiosis that originated in the Archaeplastida supergroup.

## Materials and Methods

### Chemicals

All chemicals used in this study were ACS grade or better. Biliverdin IXα was purchased from Frontier Scientific (Newark, Delaware, USA). Ferredoxin (PetF) and Ferredoxin-NADP^+^ reductase (FNR or PetH) were recombinantly expressed and purified as described elsewhere (Dammeyer *et al*., 2008; Busch *et al*., 2011). HPLC-grade acetone, formic acid and acetonitrile were purchased from VWR Chemicals (Darmstadt, Germany). Blue Sepharose™ 6 Fast Flow and Superdex™ 75 10/300 GL were purchased from GE Healthcare (Düsseldorf, Germany). Other reagents were purchased from Sigma-Aldrich (Taufkirchen, Germany).

### Plasmids and primers

Gene fragments (accession codes in Table 1) were purchased from Twist Bioscience (South San Francisco, California, USA) and Integrated DNA Technologies, Inc. (Coralville, Iowa, USA). The plasmids in Table 1 were generated via Gibson assembly (Gibson *et al*., 2009), employing primers listed in Table 2, except for pASK-IBA45(+)_*GsPEBB*. In that case, the gene was subcloned from the expression vector pCR2.1 (Eurofins Genomics Germany GmbH, Ebersberg, Germany) with *Eco*RI and *Not*I restriction sites, at the 5’ and 3’ end, respectively. *GsPEBB* was codon-optimizedized for *E. coli* using the algorithm provided by the Eurofins Genomics website, whereas *GsPEBA* was codon-harmonized using the CHARMING algorithm (http://www.codons.org/codons.html) (Wright *et al*., 2022).

**Table 1.** Plasmids.

| Plasmid | Features | Accession code |
| --- | --- | --- |
| pET28a_GsPEBA | pET-28a(+) derivative carrying the codon-harmonized *PEBA* gene sequence from *Galdieria sulphuraria* (from L70 to S302); N-terminal His<sub>6</sub>-tag | Gasu_55890 (EnsemblPlants) |
| pASK-IBA45(+)_GsPEBB | pASK-IBA45(+) derivative carrying the codon-optimized *PEBB* gene sequence from *Galdieria sulphuraria* (from N64 to T306); N-terminal Strep-tag II | Gasu_58250 (EnsemblPlants) |
| pGEX-6P-1_CmPCYA | pGEX-6P-1 derivative carrying the original *PCYA* gene sequence from *Cyanidioschyzon merolae*; N-terminal GST-tag | CMG110C (EnsemblPlants) |

**Table 2.**
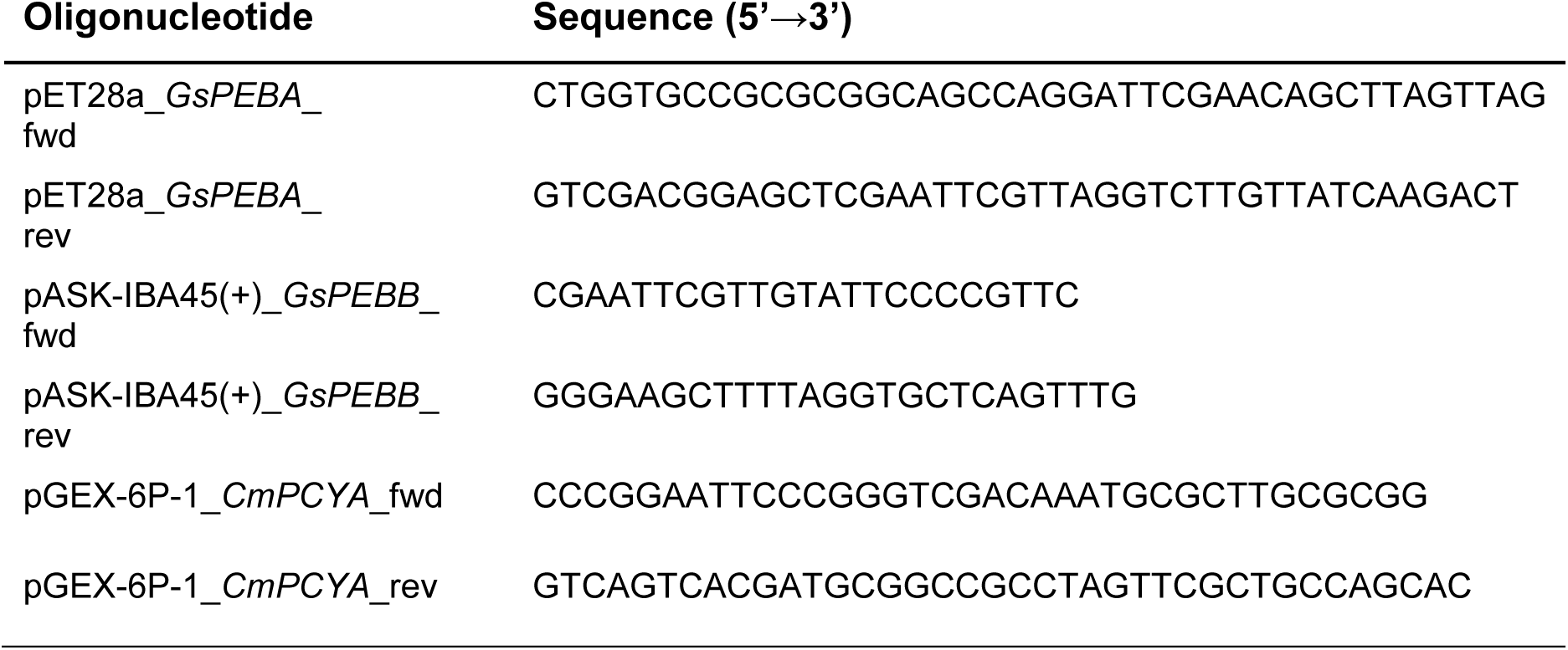
Primers.

### Production and purification of recombinant proteins

For production of recombinant His-tagged *Gs*PEBA, 3 L of LB medium supplemented with 50 µg/mL kanamycin was inoculated 1:100 with an overnight culture of *E. coli* BL21(DE3) carrying pET28a_*GsPEBA*. The cultures were grown at 37°C and 100 rpm (New Brunswick™ INNOVA^®^ 44) to an OD_600_ of 0.4 - 0.6. The temperature was decreased to 17°C and gene expression was induced by supplementing with 1 mM isopropyl-ß-thiogalactoside (IPTG). The cultures were incubated under shaking for 19 additional hours and harvested by centrifugation for 10 min at 17000 × g and 4°C (Sorvall LYNX 6000, Rotor F9). The pellet containing His-tagged *Gs*PEBA was resuspended in His-Binding buffer (20 mM sodium phosphate pH 7.4; 500 mM NaCl) at a ratio of 3 mL buffer/g wet cell weight. After the addition of 5 µg/mL DNaseI (AppliChem GmbH, Darmstadt, Germany) and 1 mg/mL lysozyme (Sigma-Aldrich), the suspension was kept on ice for 30 min. The cells were disrupted using a microfluidizer (LM 10 Microfluidizer, Microfluidics™; 3 cycles at 15000 psi) and centrifuged for 45 min at 50000 × g and 4°C (Sorvall LYNX 6000, Rotor T29). The crude extract was loaded onto a gravity flow column containing 2 mL of TALON^®^ Superflow™ resin (Cytiva, Freiburg im Breisgau, Germany). After washing with 10 column volumes (CV) of His-binding buffer, elution was performed using 4 CV of His-elution buffer (20 mM sodium phosphate pH 7.4; 500 mM NaCl; 500 mM imidazole).

For production of recombinant Strep-tagged *Gs*PEBB, 3 L LB medium supplemented with 100 µg/mL ampicillin was inoculated 1:100 with an overnight culture of *E. coli* BL21(DE3) carrying pASK-IBA45(+)_*GsPEBB*. Cells were grown at 37°C and 100 rpm (New Brunswick™ INNOVA^®^ 44) to an OD_600_ of 0.4 - 0.6. The temperature was decreased to 17°C and gene expression was induced supplementing 200 ng/mL anhydrotetracycline (AHT). The cultures were incubated under shaking for 19 additional hours and harvested by centrifugation for 10 min at 17000 × g and 4°C (Sorvall LYNX 6000, Rotor F9). The pellet containing Strep-*Gs*PEBB was resuspended in Strep-Binding buffer (100 mM Tris-HCl pH 8; 300 mM NaCl; 1 mM EDTA) at a ratio of 3 mL buffer/g wet cell weight. After the addition of a spatula tip of DNaseI and lysozyme, the suspension was kept on ice for 30 min. The cells were disrupted, centrifuged using the same conditions as for *Gs*PEBA (see above). The crude extract containing Strep-*Gs*PEBB was loaded onto a gravity flow column containing 3 mL of Strep-Tactin^®^ Sepharose^®^ (IBA Lifesciences GmbH, Göttingen, Germany). After washing with 10 CV of Strep-Binding buffer, elution was performed using 4 CV of Strep-Elution buffer (100 mM Tris-HCl pH 8; 300 mM NaCl; 1 mM EDTA; 2.5 mM Desthiobiotin).

For the production of recombinant GST-tagged *Cm*PCYA, 2 L of LB medium supplemented with 100 µg/mL ampicillin and 34 µg/mL chloramphenicol was inoculated 1:100 with an overnight culture of *E. coli* BL21(DE3) carrying pGEX-6P1_*CmPCYA* and pGro7, to support protein folding. Cells were grown at 37°C and 100 rpm (New Brunswick™ INNOVA^®^ 44) to an OD_600_ of 0.4 - 0.6. The temperature was decreased to 17°C and gene expression was induced by the addition of 0.05 mM IPTG. The cultures were incubated under shaking for 3 additional hours and harvested by centrifugation for 10 min at 17000 × g and 4°C (Sorvall LYNX 6000, Rotor F9). The pellet containing GST-tagged *Cm*PCYA was resuspended in phosphate-buffered saline (PBS:140 mM NaCl; 10 mM Na2HPO4; 2.7 mM KCl; 1.8 mM KH2PO4; pH 7.4) supplemented with 0.05% Triton X-100 at a ratio of 3 mL buffer/g wet cell weight. After the addition of a spatula tip of DNaseI and lysozyme, the suspension was kept on ice for 30 min. The cells were disrupted by sonication (Bandelin Sonopuls HD 2200; tip KE76; 5’, 5’’ pulses, 10’’ pauses; cycle 5/10; ≈ 50% power output) and centrifuged for 45 min at 4°C and 50000 × g (Sorvall^TM^ LYNX^TM^ 6000, rotor T29-8). The crude extract containing GST-*Cm*PCYA was loaded onto a gravity flow column containing 2 mL of Protino^®^ Glutathione Agarose 4B (Macherey-Nagel GmbH & Co. KG, Düren, Germany). After washing with 10 CV of PBS, elution was performed using 3 CV of GST-Elution buffer (50 mM Tris-HCl pH 8; 10 mM reduced Glutathione).

After elution, fractions for each protein were pooled and dialyzed overnight against TES-KCl buffer (25 mM TES/KOH pH 7.5; 100 mM KCl; 10%_v/v_ glycerol).

### Anaerobic bilin reductase assay

The anaerobic bilin reductase activity assays were conducted following the method outlined by Ledermann and colleagues with slight modifications (Ledermann *et al*., 2016). Recombinant ferredoxin PetF from the cyanophage P-SSM2 served as the electron donor at a final concentration of 1 µM. PetF was reduced using recombinant FNR from *Synechococcus* sp. PCC7002 at a final concentration of 0.01 µM (Busch *et al*., 2011). The reaction was initiated by adding an NADPH-regenerating system consisting of 65 mM glucose-6-phosphate, 8.2 µM NADP+ and 11 U/ml glucose-6-phosphate dehydrogenase. Absorption spectra were measured using an Agilent 8453 spectrophotometer. The reaction was stopped by 10-fold dilution of the reaction mix into ice cold 0.1% trifluoroacetic acid (TFA; v/v). Products were then isolated via solid phase extraction using Sep-Pak^®^ C18 Plus Light cartridges (Waters, Milford, Massachusetts, USA) and freeze-dried using an Alpha 2-4 LSC plus lyophilizer (Martin Christ GmbH, Osterode, Germany) prior to HPLC analysis.

### HPLC analyses

Isolated assay products were analyzed using an Agilent 1100 series HPLC system equipped with a Luna^®^ 5 µm reversed phase C18-column (Phenomenex, Torrance, California, USA) and a diode-array detector. The mobile phase consisted of 50% (v/v) acetone and 50% (v/v) aqueous 20 mM formic acid, flowing at 0.6 mL/min. Reaction products were identified by comparing their retention times with known standards and through full-spectrum analysis of the elution peaks.

### *Galdieria sulphuraria* cultivation and cell disruption

Growth medium (composition listed in Supporting Table S1) was inoculated with a single colony of *G. sulphuraria* 074W and incubated under continuous light (50 μE⋅m^−2^⋅s^−1^) at 37°C and 120 rpm (Innova^®^ 44R, New Brunswick Scientific). After reaching the exponential phase, cells were harvested by centrifugation at 3100 × g for 10 min (Eppendorf 5810 R, Rotor A-4-62) and stored at -20°C.

Approximately 7 g cells were thawed and washed three times each with H_2_O and then with Extraction buffer (50 mM HEPES; 5 mM EDTA; 10% (v/v) Glycerol; pH 7.3). Cells were resuspended in 40 mL Extraction buffer and lysed via two passages through a French^®^ Pressure Cell Press (Thermo Fisher FA-078) at 10000 PSI. The lysate was centrifuged (50000 × g, 30 min, 4 °C) to remove cell debris, and the supernatant was collected for further processing.

### *Galdieria sulphuraria* protein fraction enrichment

(NH_4_)_2_SO_4_ was finely ground using a mortar and pestle and then added to the lysate to achieve a 30% saturating concentration. The solution was stirred at 4°C for 1 h and subsequently centrifuged (50000 × g, 1 h, 4°C, Sorvall LYNX 6000, Rotor T29). The pellet was discarded, and (NH_4_)_2_SO_4_ was added to a saturating concentration of 45%. The solution was again stirred at 4°C for 1 h and centrifuged (50000 × g, 30 min, 4 °C). The pellet was dissolved in 20 mL cold Assay buffer (25 mM HEPES; 1 mM MgCl2; 10% glycerol (v/v); pH 7.3) and dialyzed at 4°C overnight against Assay buffer to remove (NH_4_)_2_SO_4_. Affinity chromatography was performed using an ÄKTA pure™ chromatography system (GE Healthcare) equipped with a Blue Sepharose™ 6 Fast Flow column (GE Healthcare) pre-equilibrated with Assay buffer. The dialyzed protein solution obtained after the last ammonium sulfate precipitation was filtered using a Phenex™-PTFE 0.45 μm syringe filter (Phenomenex) and manually loaded into the ÄKTA system with a Superloop™ (GE Healthcare). Chromatography was performed at 4°C at a flow rate of 1 mL/min. Protein elution was achieved using Elution buffer (25 mM HEPES; 1 mM MgCl2; 10% glycerol (v/v); 1 M NaCl; pH 7.3), and 5 mL fractions were collected. (NH_4_)_2_SO_4_ was added to the appropriate elution fraction at a saturating concentration of 70%, and the solution was stirred at 4°C for 1 h and then subsequently centrifuged (50000 x g, 30 min, 4 °C). The precipitated proteins were dissolved in 5 mL cold Assay buffer to perform size exclusion chromatography (SEC). Chromatography was performed on the ÄKTA™ pure 25 system equipped with a Superdex™ 75 10/300 GL column (GE Healthcare) pre-equilibrated with Assay buffer. SEC was performed at 4°C at a flow rate of 1 mL/min, and 2 mL fractions were collected. Fractions containing the desired proteins were pooled and concentrated using Amicon^®^ Ultra 4 mL Centrifugal Filters with a MWCO of 10 kDa (Merck KGaA, Darmstadt, Germany).

### Isomerase activity assay

To assess PEB:PCB isomerase activity, the purified and concentrated protein fraction was incubated with 14 μM PEB for 80 min at 30°C. Isomerase activity was monitored via UV-Vis spectroscopy using an Agilent 8453 spectrophotometer. The initial spectrum was recorded in the absence of PEB, the following one immediately after the addition of PEB and, subsequent spectra were recorded every 10 min thereafter. Following the activity assay, the reaction products were prepared for HPLC as described for the anaerobic bilin reductase assay products.

### Mass spectrometry analysis

Proteins from the isomerase-active enriched protein fraction were precipitated in 80% acetone, trypsin-digested in solution, desalted and analyzed on a HPLC-MS/MS system (Eksigent nanoLC 425 coupled to a TripleTOF 6600, Sciex, Darmstadt, Germany) as described (Hammel *et al*., 2018). Data were analyzed for peptide matching and protein inference with the MaxQuant software (v.2.6.2.0), using the *Galdieria sulphuraria* proteome from the uniprot.org repository (UP000030680, 18^th^ March 2025). Standard settings were used, except that mis-cleavages were set to a maximum of 3 and minimum peptide length to 6 amino acids.

### Phylogenetic analysis

For phylogenetic analysis of FDBRs, a multiple sequence alignment was constructed using MAFFT v7.450 (Katoh & Standley, 2014) with the command-line settings --genafpair -- maxiterate 16 --clustalout –reorder. For maximum-likelihood phylogenetic analysis, the resulting alignment was processed with an in-house script to remove positions having ≥5% gaps. Phylogenies were inferred with PhyML-3.1 (Guindon *et al*., 2010) with 100 bootstraps, using the command-line settings -m WAG -d aa -s SPR -a e -c 4 -v e -o tlr -b 100. Statistical robustness was assessed using the transfer bootstrap expectation (TBE) as implemented in booster (Lemoine *et al*., 2018). The final alignment used for tree inference contained 250 sequences, with 199 characters remaining after removal of gap-enriched columns.

## Results

All functional bilins originate through oxygenolytic cleavage of the cyclic tetrapyrrole heme (Ref HO review). In photosynthetic eukaryotes, this biosynthesis occurs in the plastid and the responsible enzymes are usually post-translationally imported into the compartment. Here, heme is first converted to BV by HO and then further reduced by FDBRs to the various bilins used in PBP and/or phytochromes. Both *Galdieria* spp. and *C. merolae* encode at least one HO in their nuclear genomes. By analogy to the data from *Gracilariopsis lemaneiformis* (Jin *et al*., 2018), we assume that the HOs in the two red algae examined here are also active in the chloroplast and synthesize BV.

### Phylogenetic analysis of rhodophyte FDBRs

Biosynthesis of PCB in *G. sulphuraria* has been proposed to proceed via synthesis of PEB and subsequent action of a putative bilin isomerase (Beale & Cornejo, 1991a, 1991b, 1991c). Subsequent examination of genomes from *Galdieria* spp. using BLAST searches and phylogenetic analyses has confirmed the presence of PEBA and PEBB and the absence of PCYA (Frankenberg *et al*., 2001; Rockwell & Lagarias, 2017), consistent with this hypothesis. However, other models also seemed possible. For example, these sequences might have been mis-assigned; other FDBRs have been identified in recent years (Chen *et al*., 2012; Rockwell *et al*., 2023; Miyake *et al*., 2025), and genomic data have become available for additional early-branching rhodophytes (Cho *et al*., 2023; Park *et al*., 2023). It has also become clear that phylogenetic analysis alone cannot reliably predict FDBR regiospecificity, as shown by the variable reaction products generated by HY2 proteins (Kohchi *et al*., 2001; Fischer *et al*., 2005; Rockwell *et al*., 2017; Frascogna *et al*., 2023) and by bacterial pre-PcyA proteins (Rockwell *et al*., 2023). We therefore began by addressing these questions.

We began by updating a recent phylogenetic analysis of FDBRs to incorporate an expanded focus on cyanobacterial sequences and on rhodophyte sequences (Rockwell *et al*., 2023). Condensed results are presented in Figure 1, with the full tree presented in Figures S2-S3 Both *G. sulphuraria* FDBRs were recovered within clades also containing sequences from other *Galdieria* spp. and from strain UTEX2393 (NCBI accession GCA_019693505.1), considered to be *Cyanidium caldiarum*. These clades themselves lay within larger branches also including sequences from other rhodophytes and from cryptophytes (Figure 1 & Figure S2). The larger PEBA clade was recovered as sister to a clade of cyanobacterial PebA sequences, whereas that for PEBB was recovered as a clade within a paraphyletic grade of cyanobacterial PebB sequences. By contrast, *C. merolae* PCYA was recovered in a clade also containing PCYA from *C. yangmingshanensis* and from a single-cell genome assigned to *C. caldiarum* (NCBI accession GCA_026184775.1). This clade also contained cryptophyte PCYA sequences (Figure 1 & Figure S3) and was part of a larger clade of PcyA/PCYA sequences from cyanobacteria and glaucophyte algae. This analysis thus demonstrates that *Gs*PEBA and *Gs*PEBB are indeed correctly assigned as PEBA and PEBB enzymes.

**Figure 1.**
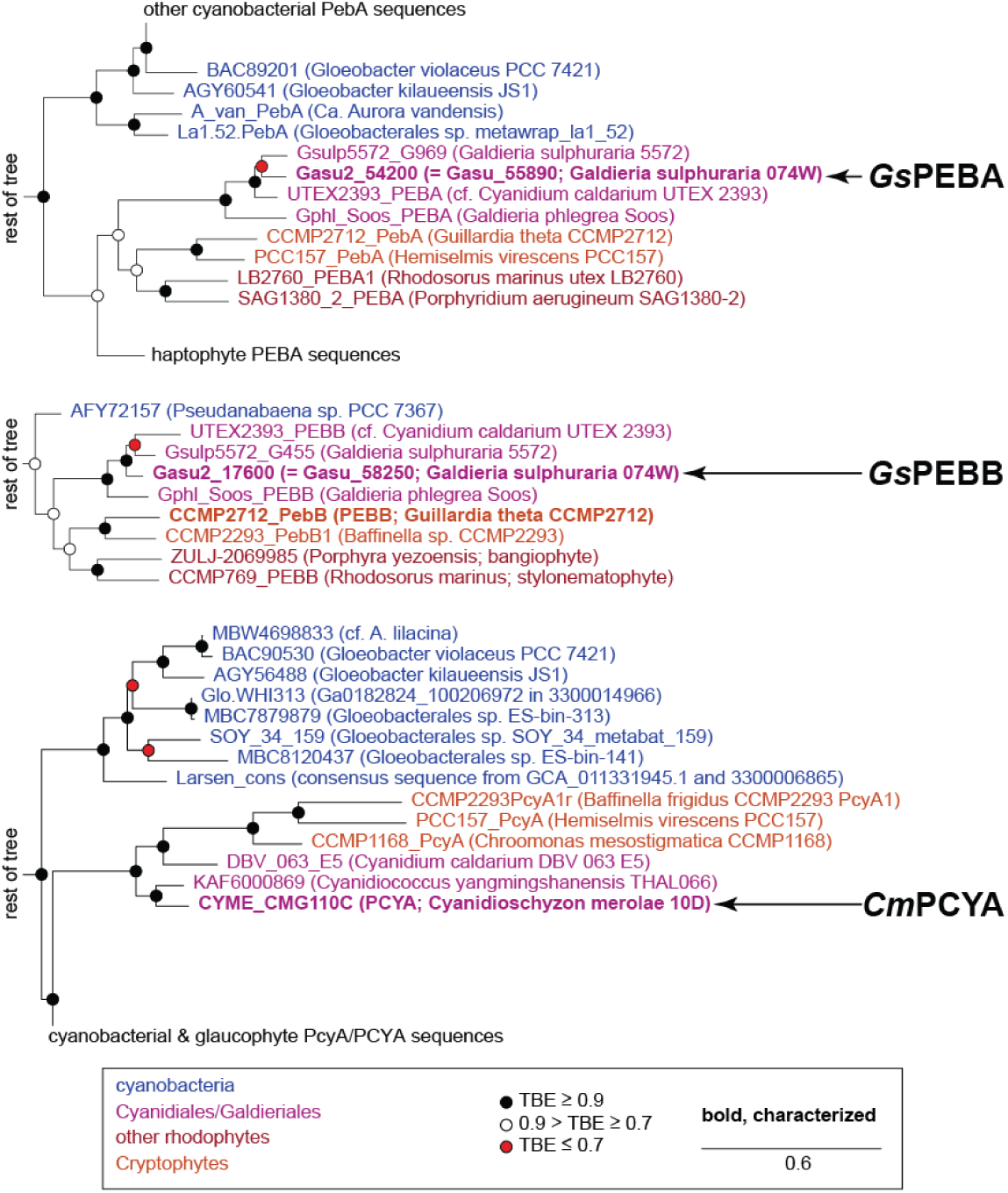
Phylogenetic analysis of FDBRs. Detailed views are shown for phylogenetic analysis of PebA/PEBA proteins (top), PebB/PEBB proteins (middle), and PcyA/PCYA proteins (bottom). The full tree is presented in Figures S2-S3. Support was evaluated using the transfer bootstrap expectation value (TBE) with 100 bootstraps as described in the Methods. Experimentally characterized proteins are indicated in **boldface**. Proteins characterized in this study are highlighted (*Gs*PEBA, *Gs*PEBB, and *Cm*PCYA).

### Biochemical characterization of FDBRs from early-branching rhodophytes

We next considered the possibility that FDBRs from *Galdieria* spp. (and those from other rhodophytes lacking PCYA) might have alternative regiospecificities allowing synthesis of PCB. Such changes in reaction products are not unknown. For example, the phage enzyme PcyX evolved from bacterial PcyA enzymes but catalyzes the synthesis of PEB rather than PCB (Ledermann *et al*., 2018). Likewise, HY2 from streptophyte algae synthesizes PCB even though HY2 from land plants instead synthesizes PΦB; this seemed particularly relevant, given the relatively close relationship between PebB/PEBB proteins and HY2 proteins (Frankenberg *et al*., 2001; Rockwell & Lagarias, 2017); also see Figure S2). We therefore characterized the *in vitro* behavior of three FDBRs: *Gs*PEBA and *Gs*PEBB from *G. sulphuraria*, and *Cm*PCYA from *C. merolae*.

We began by characterizing the reaction of His-*Gs*PEBA and BV, its expected substrate, in equimolar amounts (Figure 2A).

**Figure 2.**
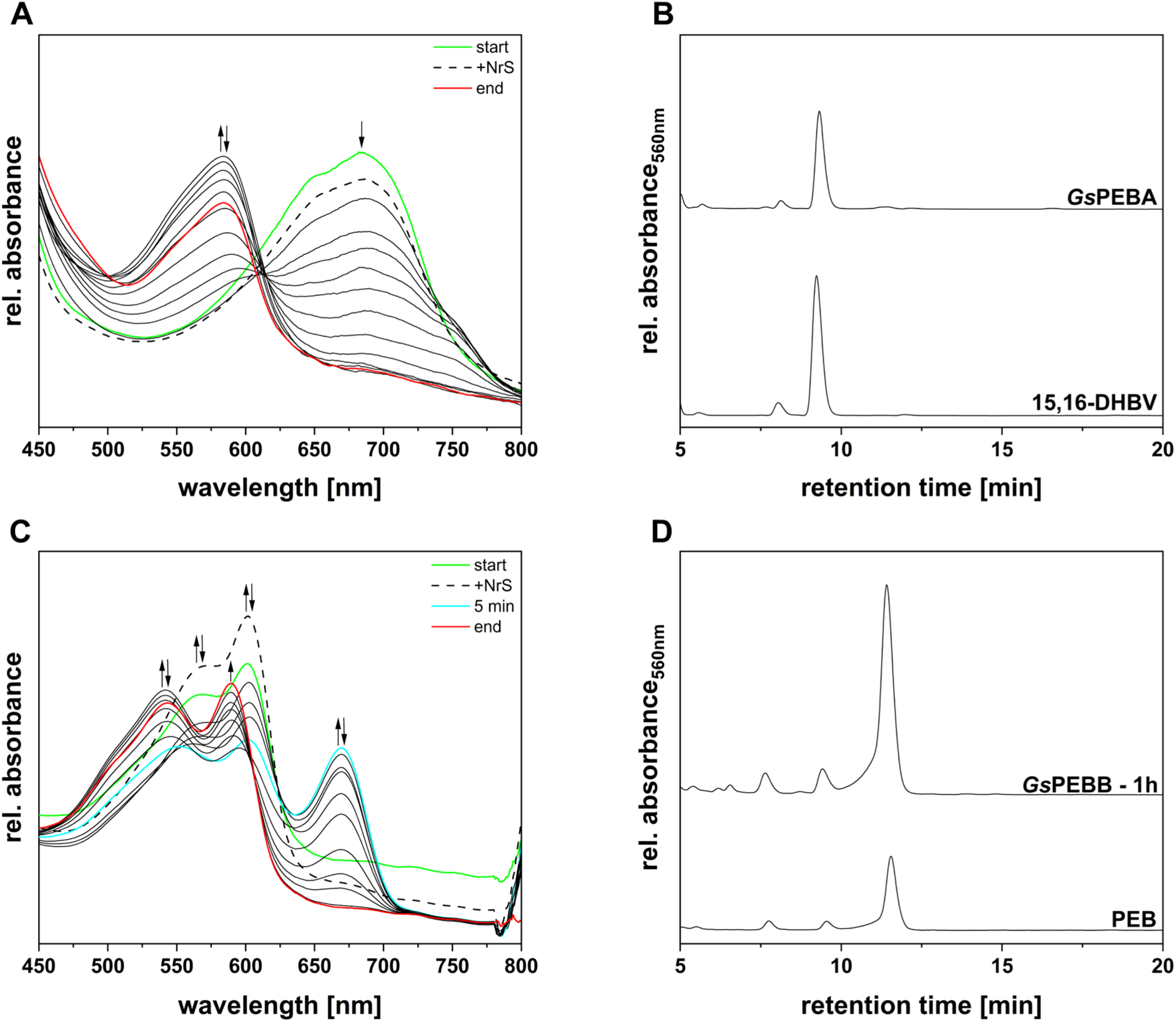
Investigation of the activity of recombinant *Gs*PEBA and *Gs*PEBB and identification of reaction products. (A) Spectra of an anaerobic bilin reductase activity assay using recombinant *Gs*PEBA and BV as the substrate. The total reaction time was 20 minutes, with spectra recorded at 30 s intervals. For ease of understanding, only the most relevant spectra are displayed. The arrows indicate the progression of absorbance during the reaction. The green spectrum corresponds to the “binding spectrum”, recorded upon incubation of BV with *Gs*PEBA. The dashed line represents the first spectrum recorded after initiating the reaction via the addition of the NrS. The spectra recorded during the reaction are shown as solid black lines, while the final spectrum is colored in red. A 40-points Savitzky-Golay filter was applied to smooth the curves. (B) HPLC analysis of the reaction products (*Gs*PEBA). The products were separated on a reversed-phase 5 μm C18 Luna column (Phenomenex), with a mobile phase consisting of 50% acetone (v/v) and 50% 20 mM formic acid (v/v), at a flow rate of 0.6 mL/min. Absorbance was monitored continuously at 560 nm. 15,16-DHBV refers to the 15,16-dihydrobiliverdin standard. (C) Spectra of an anaerobic bilin reductase activity assay using recombinant *Gs*PEBB and 15,16-DHBV as the substrate. The total reaction time was 60 minutes, with spectra recorded at 30 s intervals. For ease of understanding, only the most relevant spectra are displayed. The arrows indicate the progression of absorbance during the reaction. The green spectrum corresponds to the “binding spectrum”, recorded upon incubation of 15,16-DHBV with *Gs*PEBB. The dashed line represents the first spectrum recorded after initiating the reaction via the addition of the NrS. The spectra recorded during the reaction are shown as solid black lines, the cyan spectrum represents the product formed after 5 min, while the final spectrum is colored in red. A 40-points Savitzky-Golay filter was applied to smooth the curves. (D) HPLC analysis of the reaction products (*Gs*PEBB – 1h). The products were separated on a reversed-phase 5 μm C18 Luna column (Phenomenex), with a mobile phase consisting of 50% acetone (v/v) and 50% 20 mM formic acid (v/v), at a flow rate of 0.6 mL/min. Absorbance was monitored continuously at 560 nm. PEB refers to the phycoerythrobilin standard.

The reductase and the substrate formed a complex with an absorbance maximum at ∼ 685 nm (Figure 2A: start). Following the addition of the NADPH-regenerating system (NrS) used to generate a pool of reduced ferredoxin, a modest decrease in absorbance of the complex peak could be observed (Figure 2A: +NrS). Ultimately, *Gs*PEBA revealed activity, with the ∼ 585 nm absorbing product being accumulated within 12 minutes and subsequently undergoing non-specific degradation over time (Figure 2A: end). HPLC analysis confirmed *Gs*PEBA is indeed catalyzing the reduction of BV to 15,16-DHBV, akin to a canonical PebA (Figure 2B: *Gs*PEBA).

We next examined *Gs*PEBB, which would be expected to catalyze conversion of 15,16-DHBV to PEB. 15,16-DHBV binding to *Gs*PEBB was demonstrated by a double absorbance peak at ~ 565 nm and ∼ 600 nm (Figure 2C: start). Upon initiating the reaction, a slight increase in the absorbance of both peaks was noted, attributed to residual *Gs*PEBB:15,16-DHBV in the NrS-injection syringe (Figure 2C: +NrS). Within 5 min from the start, in addition to a decrease in the absorbance of the initial complex, a new peak at ∼ 670 nm emerged (Figure 2C: 5 min). Over time, the absorbance of these peaks decreased, ultimately resulting in a product absorbing at ~ 545 nm and ∼ 590 nm (Figure 2C: end). HPLC analysis of the final product confirmed the presence of predominantly 3(*Z*)-PEB, alongside minor traces of 3(*E*)-PEB and residual 15,16-DHBV (9.4 min product). These results demonstrate the expected activity for *Gs*PEBB and rule out any isomerizing activity of *Gs*PEBB (Figure 2D: *Gs*PEBB - 60 min).

We next considered two possibilities: first, that *Gs*PEBB might synthesize PCB from BV directly; and, second, that the two FDBRs might produce PCB when acting together. The latter hypothesis is consistent with the presence of PEB:PCB isomerase activity in a *G. sulphuraria* enriched protein fraction with MW > 60 kDa, matching the size of a two-FDBR complex. We therefore examined the reaction(s) occurring when *Gs*PEBB and *Gs*PEBA were combined in a two-enzyme activity assay (Figure 3).

**Figure 3.**
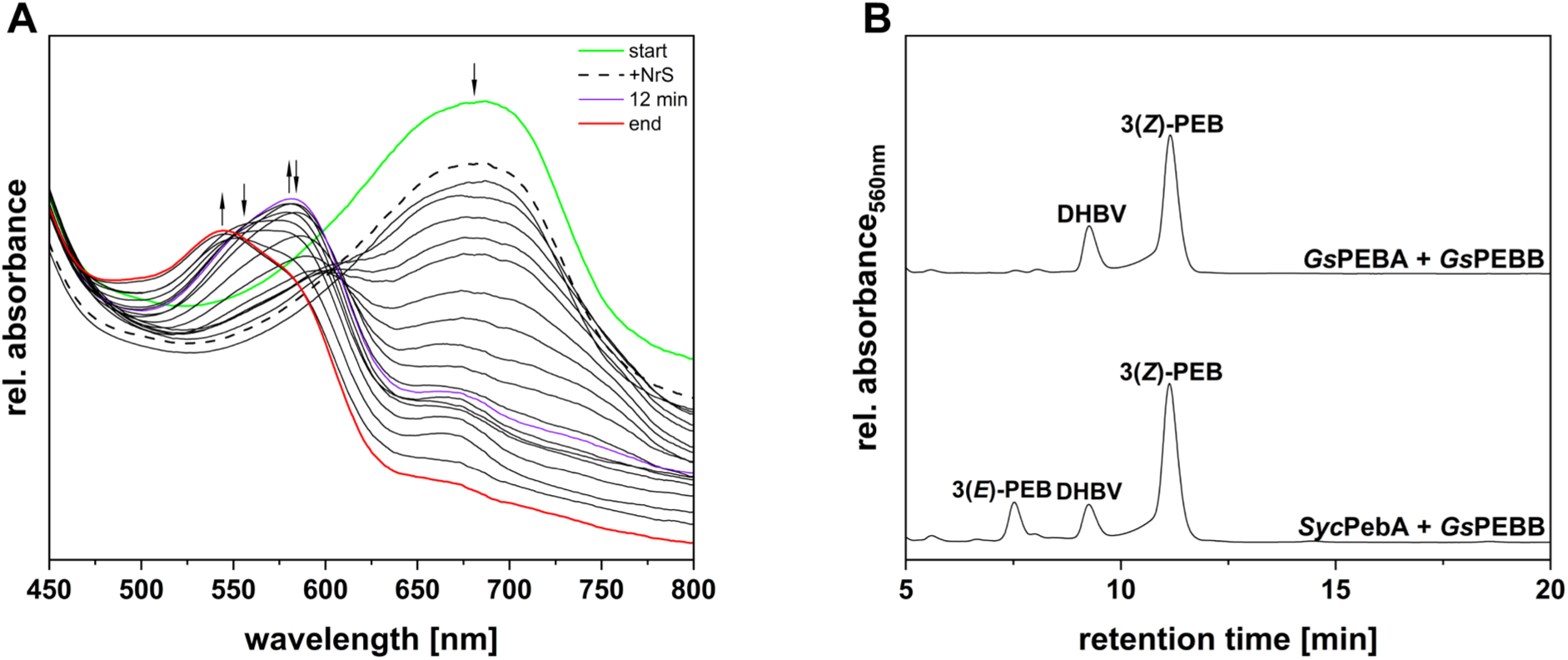
Coupled activity assay of recombinant *Gs*PEBA and *Gs*PEBB and identification of reaction products. (A) Spectra of an anaerobic bilin reductase activity assay using both recombinant *Gs*PEBA and *Gs*PEBB in a coupled approach, with BV as the substrate. The total reaction time was 40 minutes, with spectra recorded at 30 s intervals. For ease of understanding, only the most relevant spectra are displayed. The arrows indicate the progression of absorbance during the reaction. The green spectrum corresponds to the “binding spectrum”, recorded upon incubation of BV with *Gs*PEBA and *Gs*PEBB. The dashed line represents the first spectrum recorded after initiating the reaction via the addition of the NrS. The spectra recorded during the reaction are shown as solid black lines, the violet spectrum represents the product formed after 12 min, while the final spectrum is colored in red. A 60-points Savitzky-Golay filter was applied to smooth the curves. (B) HPLC analysis of the reaction products (*Gs*PEBA + *Gs*PEBB). The products were separated on a reversed-phase 5 μm C18 Luna column (Phenomenex), with a mobile phase consisting of 50% acetone (v/v) and 50% 20 mM formic acid (v/v), at a flow rate of 0.6 mL/min. Absorbance was monitored continuously at 560 nm. *Syc*PebA + *Gs*PEBB refers to the products of a coupled assay employing the PebA of *Synechococcus* sp. WH8020 and *Gs*PEBB, with BV as the substrate.

Both enzymes were added to the reaction mix from the start, in equimolar amounts, and BV was used as substrate. The spectrum obtained following the addition of the enzymes and substrate displayed a peak at ∼ 680 nm (Figure 3A: start), which decreased in absorbance after the NrS was added (Figure 3A: +NrS). After the initial formation of a product with an absorbance at ∼ 580 nm (Figure 3A: 12 min), the enzymes ultimately converted the substrate to a ∼ 545 nm absorbing product (Figure 3A: end). HPLC analysis of the final product revealed the presence of 15,16-DHBV and PEB (Figure 3B: *Gs*PEBA + *Gs*PEBB), mirroring the outcome of an equivalent reaction using PebA from *Synechococcus* sp. WH 8020 (*Syc*PebA) and *Gs*PEBB (Figure 3B: *Syc*PebA + *Gs*PEBB). This result rules out these alternative hypotheses and reaffirms the need for an isomerase activity in *G. sulphuraria*.

To finally exclude any other non-specific reaction, an additional *Gs*PEBB single activity assay was carried out using BV as the substrate. This assay confirmed that this reductase, like all other PebBs, exclusively accommodates 15,16-DHBV and does not exhibit broad substrate specificity (data not shown).

We next examined PCB biosynthesis in *Cyanidioschyzon merolae*, a close relative of *G. sulphuraria* that possesses *PCYA* (see above). *Cm*PCYA would be expected to carry out direct, 4-electron reduction of BV to PCB. Of several expression conditions, only the one in which the expression of pGEX-6P-1_*CmPCYA* was coupled with pGro7, to support protein folding, and the amount of the inducer and the incubation time were reduced, proved to be successful in obtaining an active enzyme.

Characterization of *Cm*PCYA confirmed that it is a *bona fide* PCYA. The reductase and the substrate formed a complex with an absorbance maximum at ∼ 660 nm (Figure 4A: start). Following the addition of the NrS, a modest decrease in absorbance of the complex peak could be observed (Figure 4A: +NrS). Ultimately, *Cm*PCYA-catalyzed BV reduction led to the formation of a product absorbing at ∼ 620 nm (Figure 4A: end). HPLC analysis confirmed *Cm*PCYA is consistent with its classification, catalyzing the reduction of BV to PCB (Figure 4B: *Cm*PCYA). Furthermore, the pigment eluting at ∼ 17 min potentially indicated the presence of 18^1^,18^2^-DHBV as the intermediate, as has been observed for cyanobacterial PcyA (Frankenberg & Lagarias, 2003a; Miyake *et al*., 2020). These results thus demonstrate that the rarely occurring rhodophyte PCYA sequences can indeed carry out direct PCB synthesis.

**Figure 4.**
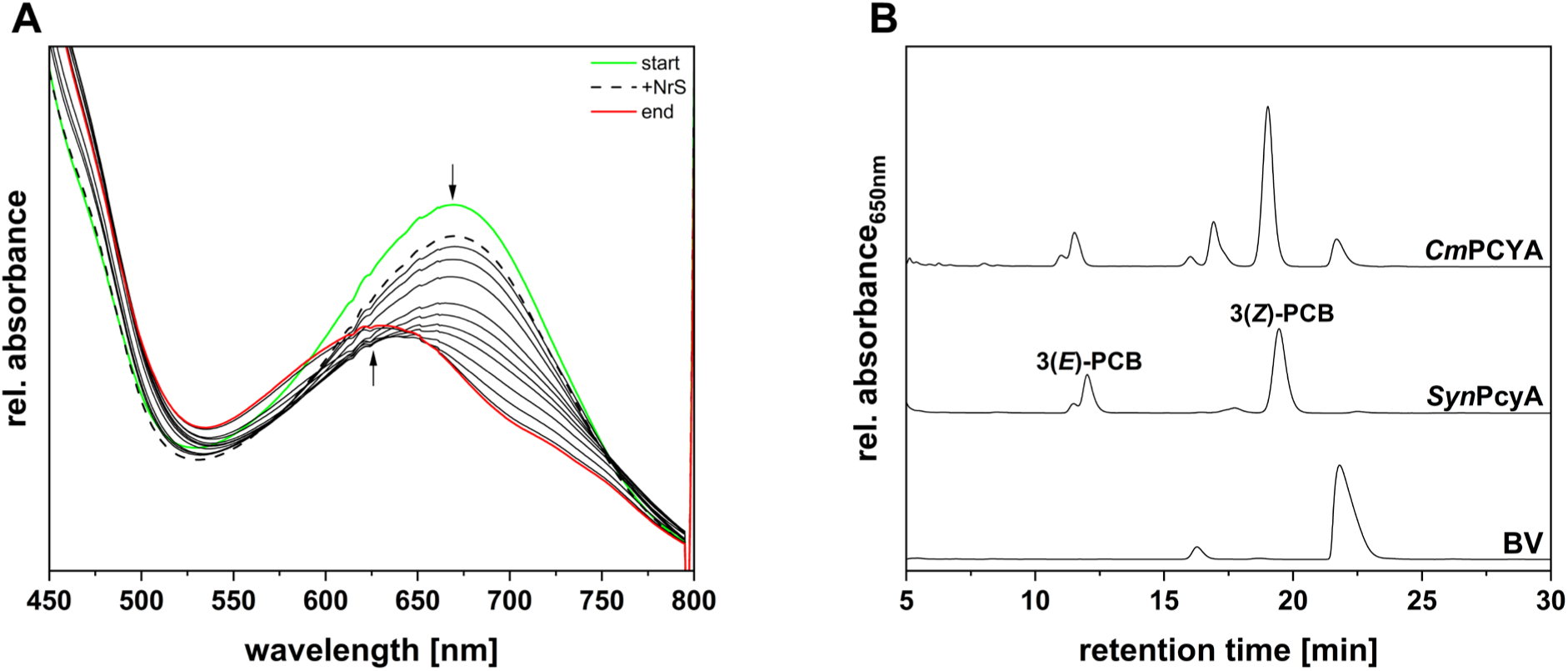
Investigation of the activity of recombinant *Cm*PCYA and identification of reaction products. (A) Spectra of an anaerobic bilin reductase activity assay using recombinant *Cm*PCYA and BV as the substrate. The total reaction time was 10 minutes, with spectra recorded at 30 s intervals. For ease of understanding, only the most relevant spectra are displayed. The arrows indicate the progression of absorbance during the reaction. The green spectrum corresponds to the “binding spectrum”, recorded upon incubation of BV with *Cm*PCYA. The dashed line represents the first spectrum recorded after initiating the reaction via the addition of the NrS. The spectra recorded during the reaction are shown as solid black lines, while the final spectrum is colored in red. A 10-points Savitzky-Golay filter was applied to smooth the curves. (B) HPLC analysis of the reaction products (*Cm*PCYA). The products were separated on a reversed-phase 5 μm C18 Luna column (Phenomenex), with a mobile phase consisting of 50% acetone (v/v) and 50% 20 mM formic acid (v/v), at a flow rate of 0.6 mL/min. Absorbance was monitored continuously at 650 nm. *Syn*PcyA refers to the product of an assay performed using *Synechocystis* sp. PCC 6803 PcyA and BV as the substrate, BV refers to the biliverdin standard.

### Isomerization of PEB to PCB

Our studies thus confirm a requirement for PEB:PCB isomerase activity in *G. sulphuraria*, consistent with the early work of Beale and Cornejo (Beale and Cornejo, 1991b). We therefore sought to confirm the presence of this activity in *G. sulphuraria*, using the protocol described in this early work. Following the mechanical disruption of the *G. sulphuraria* cells, two ammonium sulfate precipitations were carried out. After dialysis of the precipitated protein fraction from the second ammonium sulfate fractionation, Blue Sepharose affinity chromatography was performed (Figure 5A).

**Figure 5.**
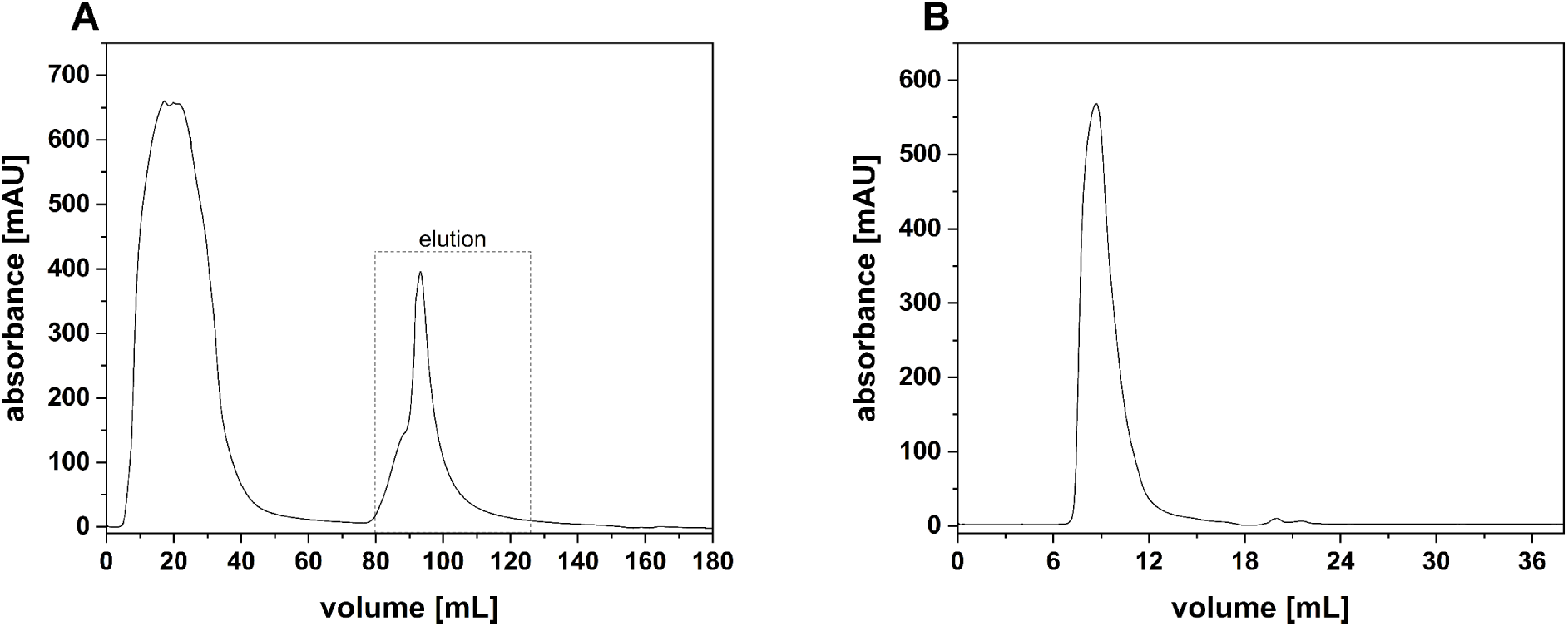
Affinity chromatography and size exclusion chromatography of a protein enriched fraction from *G. sulphuraria*. (A) Affinity chromatography was performed using the ÄKTA™ pure 25 purification system equipped with a column packed with Blue Sepharose™ 6 Fast Flow. Equilibration and washing with “Assay buffer” were followed by elution with “Elution buffer”. Absorbance was continuously measured at 280 nm. (B) Size exclusion chromatography was performed using the ÄKTA™ pure 25 purification system equipped with a Superdex™ 75 10/300 GL, equilibrated with “Assay buffer”. Absorbance was continuously measured at 280 nm.

The column material interacted with target proteins in the algal lysate, eluting at ∼ 90 mL (Figure 5A). This fraction was subsequently treated with a 75% saturating concentration of ammonium sulfate. Precipitated proteins were resuspended in assay buffer and subjected to size exclusion chromatography (Figure 5B). The previous findings of Beale & Cornejo indicated that an isolated fraction with a molecular mass greater than 60 kDa exhibited isomerase activity. Using the elution profile of albumin as reference (66 kDa, Figure S4), the fractions eluting between 7 and 12 mL were collected and employed in an isomerase activity screening. These fractions all displayed a bluish color, consistent with the possible presence of endogenous PCB but not that of the pink-colored PEB. 14 µM of purified PEB was added to 1.5 mL concentrated protein fraction, and potential conversion to PCB was monitored via spectroscopy at 30°C over the course of 80 min (Figure 6A).

**Figure 6.**
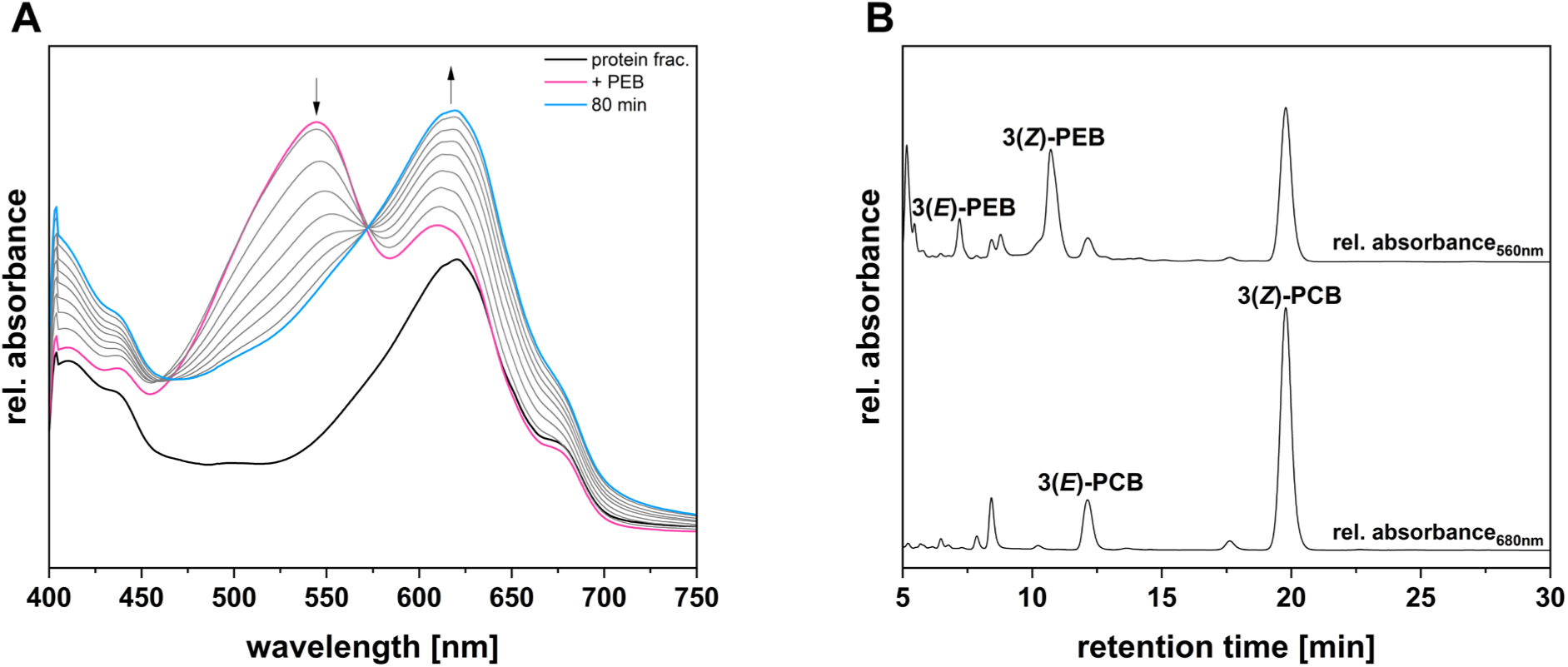
Isomerase activity assay of a *G. sulphuraria* enriched protein fraction and identification of products. (A) Isomerase activity assay of a *G. sulphuraria* enriched protein fraction. The solid black line represents the absorbance of the protein fraction (protein frac.). The enriched protein fraction was incubated with 14 μM PEB at 30°C. The pink spectrum was acquired immediately after the addition of PEB (+ PEB). The course of absorbance, indicated by the arrows, was recorded every 20 min for 80 min. The cyan spectrum represents the end spectrum, recorded at 80 min (80 min). A 10-points Savitzky-Golay filter was applied to smooth the curves. (B) HPLC analysis of the isomerase assay products. The products were separated on a reversed-phase 5 μm C18 Luna column (Phenomenex), with a mobile phase consisting of 50%^v/v^ acetone and 50%_v/v_ 20 mM formic acid, at a flow rate of 0.6 mL/min. Absorbance was monitored continuously at 560 nm and 650 nm. The upper trace indicates the chromatogram recorded at the constant DAD wavelength of 560 nm (rel. absorbance_560nm_). The lower trace indicates the chromatogram recorded at the constant DAD wavelength of 650 nm (rel. absorbance_650nm_). 3(*E*)-PEB, 3(*E*)-phycoerythrobilin; 3(*Z*)-PEB, 3(*Z*)-phycoerythrobilin; 3(*E*)-PCB, 3(*E*)-phycocyanobilin; 3(*Z*)-PCB, 3(*Z*)-phycocyanobilin.

The protein fraction initially showed an absorption peak at 620 nm, consistent with the presence of phycocyanin (PC), and shoulders at 410, 440 and 680 nm representing chlorophyll a (Figure 6A: protein frac.). Addition of pure PEB resulted in formation of a new peak at 540 nm (Figure 6A: + PEB). At the final time point, the UV-Vis spectrum displayed apparently complete conversion of the 540 nm peak to PCB, the 620 nm peak (Figure 6A: 80 min). HPLC analysis of the 80 min assay sample revealed the presence of PCB and remaining amounts of PEB (Figure 6B), whereas the double peak eluting at ∼ 8.5 min and absorbing at 560 nm could indicate the presence of 15,16-DHBV.

The dependence of this activity on an enzymatic factor was verified via negative controls (Figure 7). In the first control assay, the enriched protein fraction was heat-inactivated before the assay (Figure 7A, B). In the second control, a buffer blank was used instead of the protein fraction (Figure 7C, D). None of these control assays exhibited detectable isomerase activity. The isomerase-active protein fraction was analyzed by mass spectrometry to potentially identify the responsible protein (Supporting data set). Unfortunately, the fraction is particularly protein rich, with 356 identified proteins including ∼ 10% of unknown function. Nevertheless, the presence of phycobiliprotein-related entries is consistent with the initial blue color and 620 nm peak and may facilitate further purification steps.

**Figure 7.**
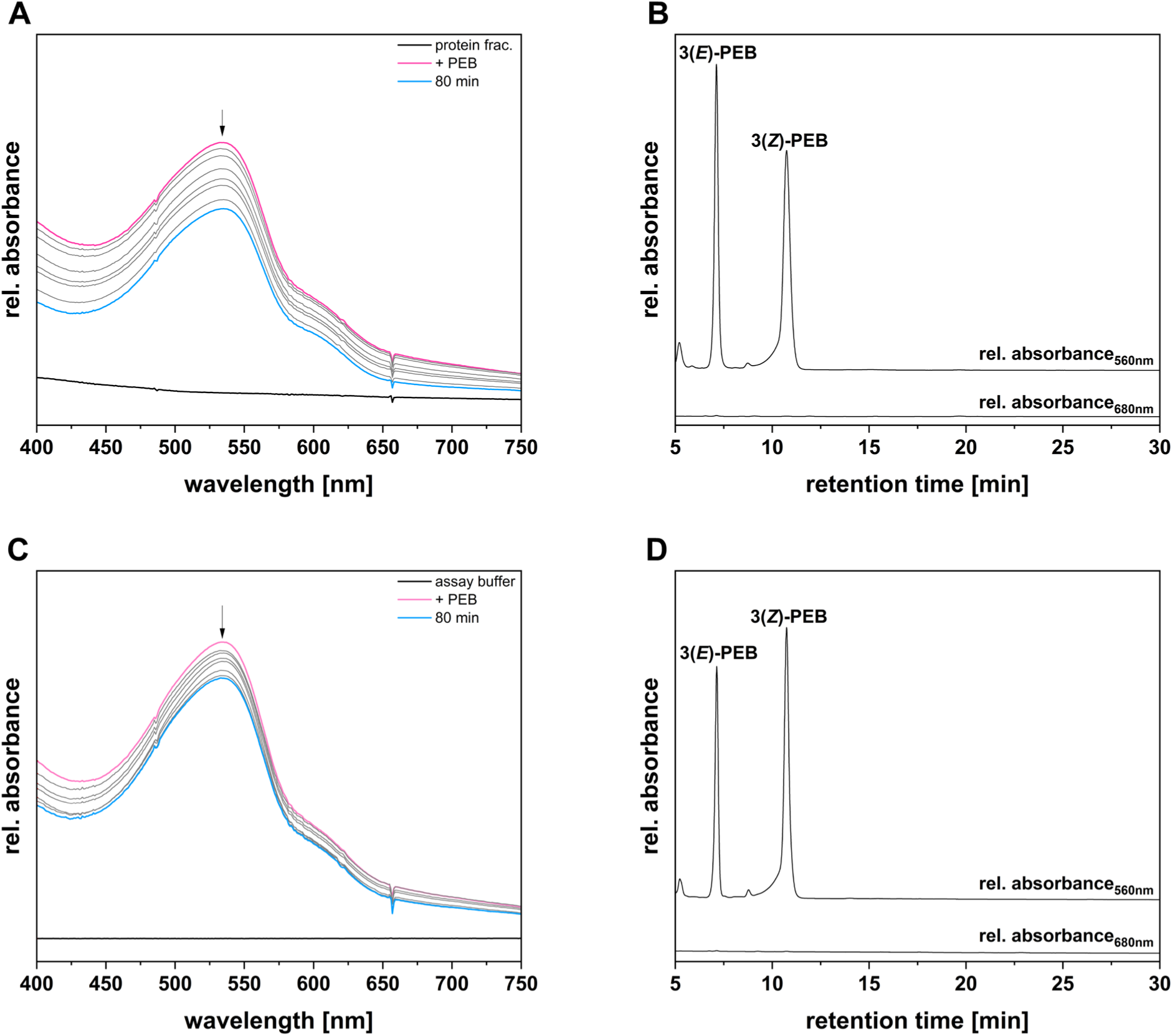
Negative controls of isomerase activity assay and identification of products. (A) Isomerase activity assay of a heat-inactivated *G. sulphuraria* enriched protein fraction. The solid black line represents the absorbance of the protein fraction subjected to heat-inactivation at 95°C for 10 min (protein frac.). The enriched protein fraction was incubated with 14 μM PEB at 30°C. The pink spectrum was acquired immediately after the addition of PEB (+ PEB). The course of absorbance, indicated by the arrow, was recorded every 20 min for 80 min. The cyan spectrum represents the end spectrum, recorded at 80 min (80 min). (B) HPLC analysis of the isomerase assay products. The products were separated on a reversed-phase 5 μm C18 Luna column (Phenomenex), with a mobile phase consisting of 50%_v/v_ acetone and 50%_v/v_ 20 mM formic acid, at a flow rate of 0.6 mL/min. Absorbance was monitored continuously at 560 nm and 650 nm. The upper trace indicates the chromatogram recorded at the constant DAD wavelength of 560 nm (rel. absorbance_560nm_). The lower trace indicates the chromatogram recorded at the constant DAD wavelength of 560 nm (rel. absorbance_650nm_). 3(*E*)-PEB, 3(*E*)-phycoerythrobilin; 3(*Z*)-PEB, 3(*Z*)-phycoerythrobilin. (C) Isomerase activity assay performed substituting the *G. sulphuraria* enriched protein fraction with “Assay buffer”. The solid black line represents the absorbance of the “Assay buffer” (buffer). The “Assay buffer” was incubated with 14 μM PEB at 30°C. The pink spectrum was acquired immediately after the addition of PEB (+ PEB). The course of absorbance, indicated by the arrow, was recorded every 20 min for 80 min. The cyan spectrum represents the end spectrum, recorded at 80 min (80 min). A 10-points Savitzky-Golay filter was applied to smooth the curves. (D) HPLC analysis of the isomerase assay products. The products were separated on a reversed-phase 5 μm C18 Luna column (Phenomenex), with a mobile phase consisting of 50%_v/v_ acetone and 50%_v/v_ 20 mM formic acid, at a flow rate of 0.6 mL/min. Absorbance was monitored continuously at 560 nm and 650 nm. The upper trace indicates the chromatogram recorded at the constant DAD wavelength of 560 nm (rel. absorbance_560nm_). The lower trace indicates the chromatogram recorded at the constant DAD wavelength of 560 nm (rel. absorbance_650nm_). 3(*E*)-PEB, 3(*E*)-phycoerythrobilin; 3(*Z*)-PEB, 3(*Z*)-phycoerythrobilin.

## Discussion

Although they are the basal clade of Rhodophyta, the Cyanidiophyceae exhibit a dark green coloration due to the absence of PE in their PBS. On the basis of the pigment composition, Cyanidiophyceae should encode only *PCYA*, responsible for synthesizing PCB to be subsequently bound to APC and PC, whereas other rhodophytes would also have *PEBA* and *PEBB* genes. This simple expectation does not hold true, providing the impetus for this study. Consistent with previous studies, our phylogenetic analysis demonstrates that rhodophyte PCYA sequences are only found in the early-branching genera *Cyanidioschyzon* and *Cyanidiococcus*, along with a sequence in one of the two available genomes assigned to *Cyanidium*. All other rhodophyte genomes examined to date have only *PEBA* and *PEBB*, despite the need for PCB chromophores in the rhodophyte PBS (Zhang *et al*., 2017; Ma *et al*., 2020). Our phylogenetic analysis also provides several other insights. First, there is an apparent discrepancy in the two available genomes classified as *Cyanidium caldarium*: strain UTEX 2393, with genome accession GCA_019693505.1, has *PEBA* and *PEBB* and resembles *Galdieria* spp., but a single-cell genome with accession GCA_026184775.1 instead has *PCYA*. It will thus be interesting to see the FDBR composition as other nuclear genomes become available for early-diverging rhodophytes. We also observe that all three rhodophyte FDBRs are associated with candidate cryptophyte orthologs (Figure 1), in contrast to the FDBRs found in other secondary algae with plastids derived from endosymbiosis with red algae (Rockwell & Lagarias, 2017; Rockwell *et al*., 2023). As in rhodophytes, cryptophytes have frequently lost PCYA. Cryptophyte PCYA proteins form a clade but are found in diverse genera that are not closely related to each other, consistent with acquisition of all three FDBRs at establishment of secondary endosymbiosis with subsequent loss. Our studies therefore indicate that the ancestral rhodophyte would also have had all three cyanobacterial FDBRs and that the loss of PCYA occurred quite early in rhodophyte evolution. Hence, even the early-branching *G. sulphuraria* has an apparent need for an isomerase to convert PEB into PCB for light harvesting. Although the two FDBRs, *Gs*PEBA and *Gs*PEBB, are classified phylogenetically according to their predicted activities, past experience has repeatedly shown that biochemical analyses are essential to confirm their final activities (Dammeyer *et al*., 2008; Rockwell *et al*., 2017; Ledermann *et al*., 2018; Frascogna *et al*., 2023). We therefore confirmed the activity of representative PCYA, PEBA, and PEBB proteins from early-branching rhodophytes. PEBA and PEBB from *G. sulphuraria* were proven both in individual and in coupled assays to be responsible for the production of only 15,16-DHBV and PEB, respectively. Therefore, production of PCB would be expected to use PEB as a precursor, because both bilins are at the same oxidation state and the reaction would thus be a simple isomerization reaction as proposed by Beale and Cornejo (Figure 8) (Beale & Cornejo, 1991b). The same pathway is likely to be present in more complex Rhodophytes (Rhodophytina) as well. In such organisms, PE is also present in the PBS (Zhang *et al*., 2017; Ma *et al*., 2020), but PCB would still be required for energy transfer from PE to PC and then APC. The presence of PE explains the existence of *PEBA* and *PEBB* in these organisms, yet no *PCYA* is encoded for direct PCB biosynthesis starting from BV, again requiring an isomerase activity.

**Figure 8.**
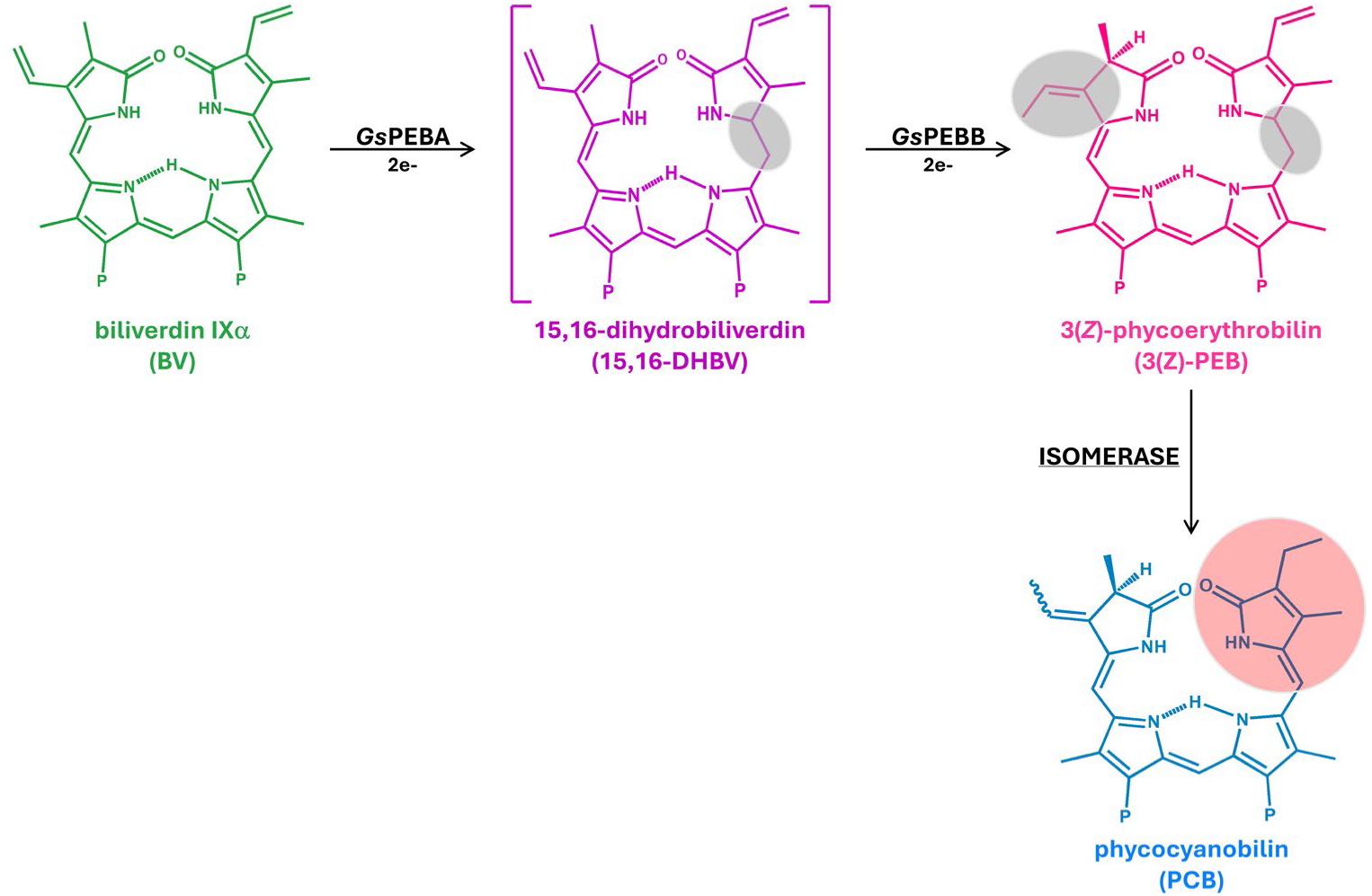
Proposed PCB biosynthesis pathway in *Galdieria sulphuraria*. *Gs*PEBA catalyzes the 2e^-^ reduction of BV to 15,16-DHBV. *Gs*PEBB subsequently reduces 15,16-DHBV to mostly 3(*Z*)-PEB, with small traces of the 3(*E*) isomer. “P” indicates the propionate side chains. BV reduction sites are highlighted in gray. 3(*Z*)-PEB is the substrate for an isomerase acting on the C18^1^=C18^2^ double bond. Isomerization site is highlighted in red.

Through fractionation of *G. sulphuraria* extracts, our research group also confirmed the isomerase activity proposed in 1991. In particular, a fraction enriched with *G. sulphuraria* protein with MW > 60 kDa was found to convert exogenous PEB to PCB. However, an attempt to identify the putative isomerase in this fraction via mass spectrometry was not successful. Almost 400 proteins were identified in the sample, of which 14 remain uncharacterized and nine are annotated as isomerases. Future work might permit further enrichment of this fraction, reducing the number of potential candidates to allow a recombinant approach for the characterization of the putative isomerase.

In *silico* analysis using bioinformatics was also performed as another potential approach to facilitate the identification of the isomerase. As previously mentioned, the Cyanidioschyzonales and the Cyanidiales, sister lineages to the Galdieriales, harbor *PCYA*. We confirmed that *Cm*PCYA is a genuine PCYA, catalyzing reduction of BV to PCB. Taking advantage of this significant distinction in the bilin biosynthesis pathway, a proteome subtraction was conducted between *G. sulphuraria* and *C. merolae*. *C. merolae* has a reduced genome (16 Mb) with approximately 4775 protein-coding genes (Nozaki *et al*., 2007). *G. sulphuraria* is slightly more complex, with an even smaller genome (13.7 Mb) but with a higher number (approximately 6622) of protein-coding genes (Schönknecht *et al*., 2013). However, the subtraction of the two proteomes was not particularly informative: approximately 4100 proteins were unique to *G. sulphuraria*, consistent with the findings from Schönknecht and colleagues, and ∼ 300 of these proteins remain uncharacterized. Additional comparison of these unique proteins with the ones identified by mass spectrometry in the enriched fraction led to tentative identification of 30 potential candidates, all encoded in the nucleus (Table S2). Nevertheless, the amount remains excessive to proceed with the characterization of each in a recombinant approach. It may instead be possible to re-engineer existing heterologous expression systems to characterize these proteins, because *E. coli* systems have already been created for biosynthesis of PEB and for expression of cyanobacteriochrome proteins that preferentially bind PCB and not PEB (Rockwell *et al*., 2012). Combining such systems might allow screening for isomerase activity, although the absence of a positive control presents an obvious concern. We also considered the possibility that PCB formation might proceed via the action of a bilin lyase-isomerase. In this model, a PEB-specific lyase-isomerase carries out isomerization of PEB to PCB and attachment to PC and APC subunits in one step. Such mechanisms are typical of cyanobacteria, where the synthesis of PUB and PVB and their respective attachment to PE and PEC are based on lyase-isomerases (Zhao *et al*., 2005, 2017; Blot *et al*., 2009; Shukla *et al*., 2012; Mahmoud *et al*., 2017; Sanfilippo *et al*., 2019; Carrigee *et al*., 2021, 2022). Although bilin lyases and lyase-isomerases remain poorly characterized in rhodophytes, BLAST searches led to the identification of the PC-related enzymes CpcE/F, CpcS and CpcT in both *C. merolae* and *G. sulphuraria* (Table S3). *G. sulphuraria* uniquely possesses CpcU, but this protein is thought to only function as a heterodimer with CpcS (Zhao *et al*., 2006, 2007; Saunée *et al*., 2008). Hence, the apparent palette of bilin lyases is quite similar in both organisms, even though only one has a need for an isomerase activity.

Future work to identify the bilin isomerase of *G. sulphuraria* may also benefit from new methodological approaches. Recently, the development of a click-chemistry PCB probe (a PCB adduct bearing a biotinylated C12-alkyne) allowed for the identification of PCB-interacting proteins by adding the probe to mammalian cells and subsequent enrichment via biotin-avidin affinity (Wilkinson *et al*., 2023). In the search for the isomerase, a similar strategy could be envisaged with the use of a PEB clickable probe. Introducing this probe to cell-free extracts or plastid isolates could potentially facilitate screening for the isomerase among the potential pool of interacting partners. However, the success of this approach is highly dependent on efficient cell breakage, which is challenging in *G. sulphuraria* due to the highly proteinaceous cell wall (Mercer *et al*., 1962; Bailey & Staehelin, 1968; Oesterhelt *et al*., 2008). Ideally, utilizing the more tractable *G. partita* (Hirooka *et al*., 2022) would provide a lysis advantage and would additionally reduce potential PEB cross-reactions. Since the absence of *PCYA* is also a common trait of Rhodophytina, the search for the isomerase could be moved to a more complex organism. However, the greater complexity and larger genomes of such organisms may only partially offset the challenges associated with easier cell lysis.

## Supporting information

Supporting data_set

## Acknowledgement

We are thankful to Andreas P.M. Weber for the gift of the *Galdieria sulphuraria* and *Cyanidioschyzon merolae* strains and J. Clark Lagarias for helpful discussion and continuous support. Furthermore, we would like to thank Frederik Sommer and Michael Schroda for the Mass Spectrometry analysis.

## Author contributions

FF, NCR, and NFD designed research; FF, JH, JM, and NCR performed research; FF, NCR, and NFD wrote the paper. All authors read and approved the final version of the manuscript.

## Data availability

Raw data are available on DataDryad: https://doi.org/10.5061/dryad.hhmgqnksx

## Supporting Information

Supporting Information may be found online in the relative section at the end of the article.

**Figure S1.** Bilin reductases overview.

**Figure S2.** Phylogenetic analysis of PebA and PebB lineages.

**Figure S3.** Phylogenetic analysis of the PcyA lineage.

**Figure S4.** Size exclusion chromatography of bovine serum albumin (BSA).

**Table S1.** *Galdieria sulphuraria* growth media and Trace elements composition.

**Table S2.** Proteins exclusively identified in the isomerase-active enriched fraction of *Galdieria sulphuraria*, absent in the proteome of *Cyanidioschyzon merolae*.

**Table S3.** Comparison of phycobiliprotein lyases in *Galdieria sulphuraria* and *Cyanidioschyzon merolae*.

**Supporting data set.** Mass spectrometry analysis of the isomerase-active protein fraction.

