## supporting Material for "Phycocyanobilin biosynthesis in *Galdieria sulphuraria* requires isomerization of phycoerythrobilin synthesized by bilin reductases"

---

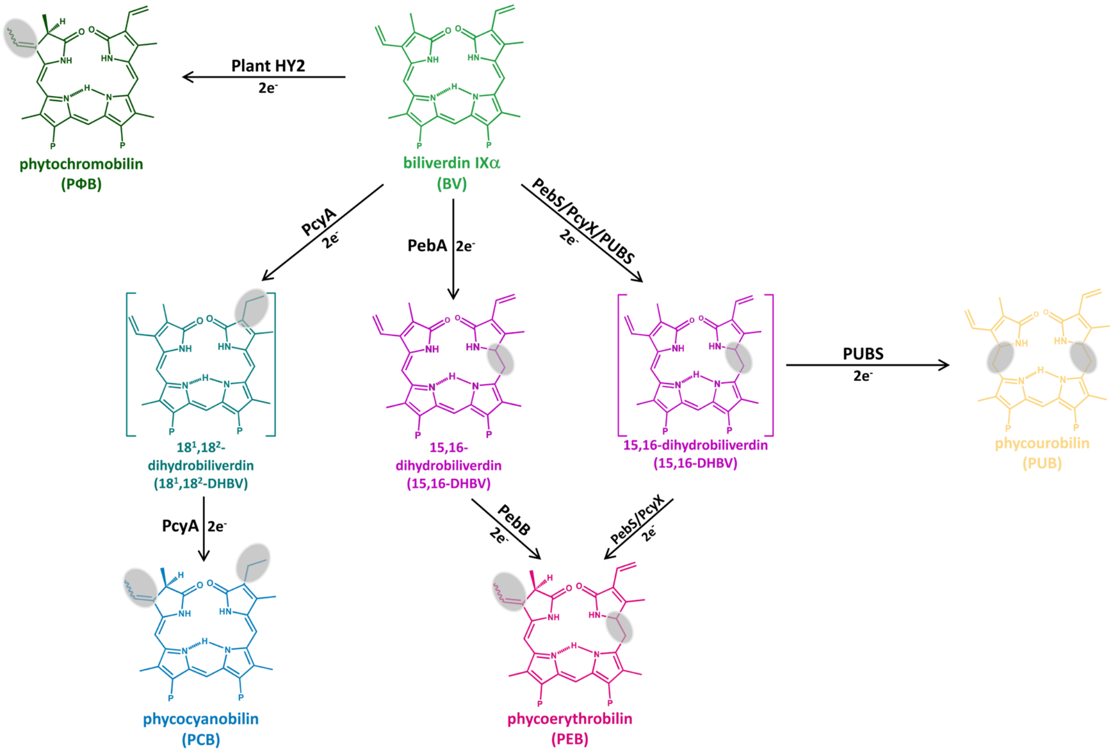


**Supporting Figure S1. Bilin reductases overview**. Reduction sites are highlighted in grey. The plant enzyme phytochromobilin synthase (HY2) is responsible for the reduction of biliverdin (BV) A-ring to yield phytochromobilin (PΦB) (Frankenberg *et al.*, 2001; Kohchi *et al.*, 2001). A second type of HY2, the one of streptophyte algae, catalyzes the reduction of BV to phycocyanobilin (PCB), either via PΦB or $18^1$,$18^2$-dihydrobiliverdin ($18^1$,$18^2$-DHBV) (Rockwell *et al.*, 2017; Frascogna *et al.*, 2023). PcyA catalyzes the $4e^-$ reduction of BV first to the intermediate $18^1$,$18^2$-DHBV and ultimately to PCB (Frankenberg & Lagarias, 2003). PebA reduces BV to 15,16-dihydrobiliverdin (15,16-DHBV), while PebB converts the latter to phycoerythrobilin (PEB) (Dammeyer & Frankenberg-Dinkel, 2006; Busch *et al.*, 2011). PebS and PcyX combines the activity of PebA and PebB in one enzyme (Dammeyer *et al.*, 2008; Ledermann *et al.*, 2018). PUBS, only found in Viridiplantae, catalyzes the $4e^-$ reduction of BV to phycourobilin, via 15,16-DHBV as intermediate (Chen *et al.*, 2012; Miyake *et al.*, 2025).

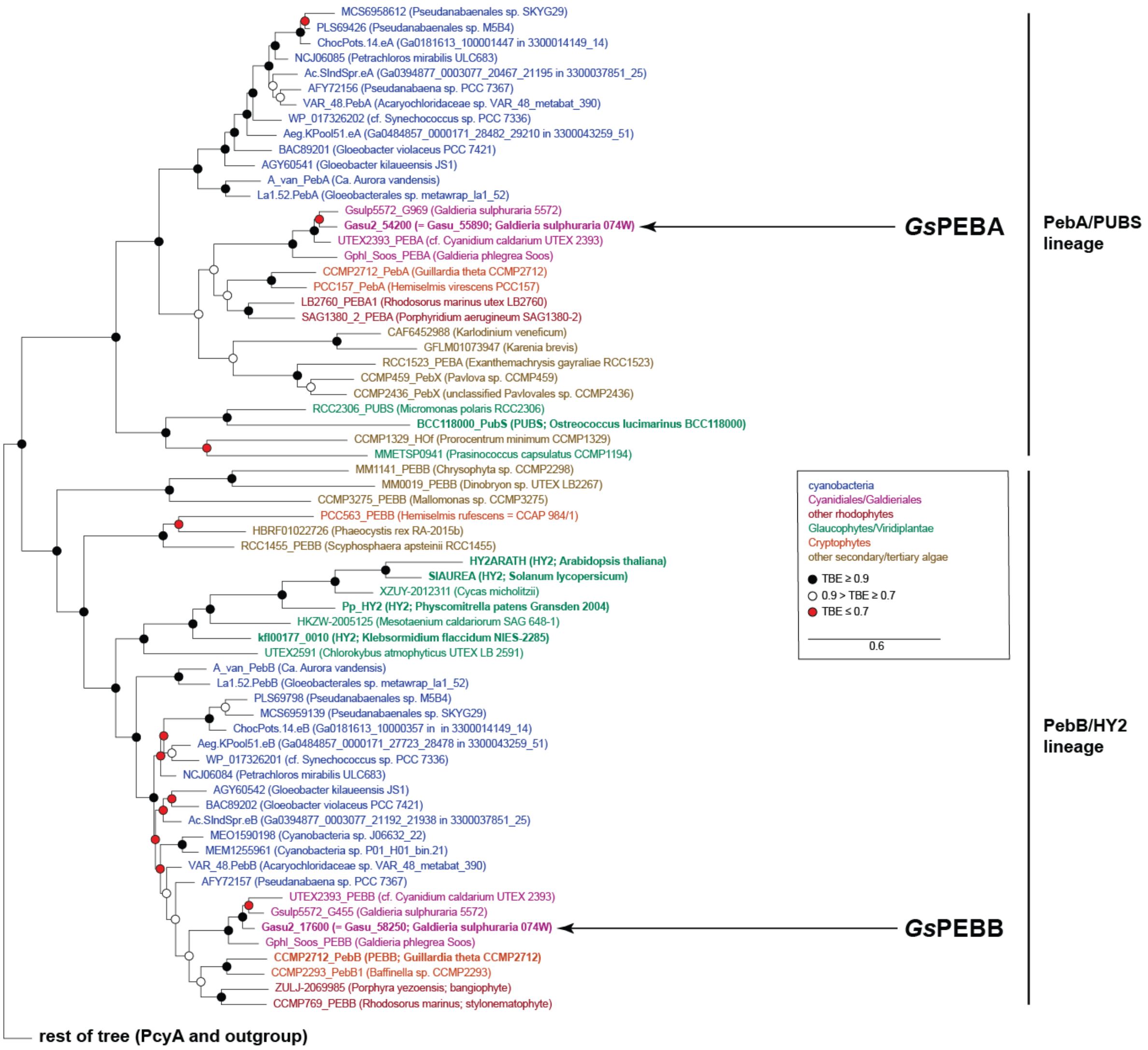


**Supporting Figure S2. Phylogenetic analysis of PebA and PebB lineages.** The section of the tree containing the PebA/PEBA/PUBS and PebB/PEBB/HY2 lineages is shown, with the rest of the tree presented in Supporting Figure S3. Details of tree construction are presented in the Methods. TBE, transfer bootstrap expectation.

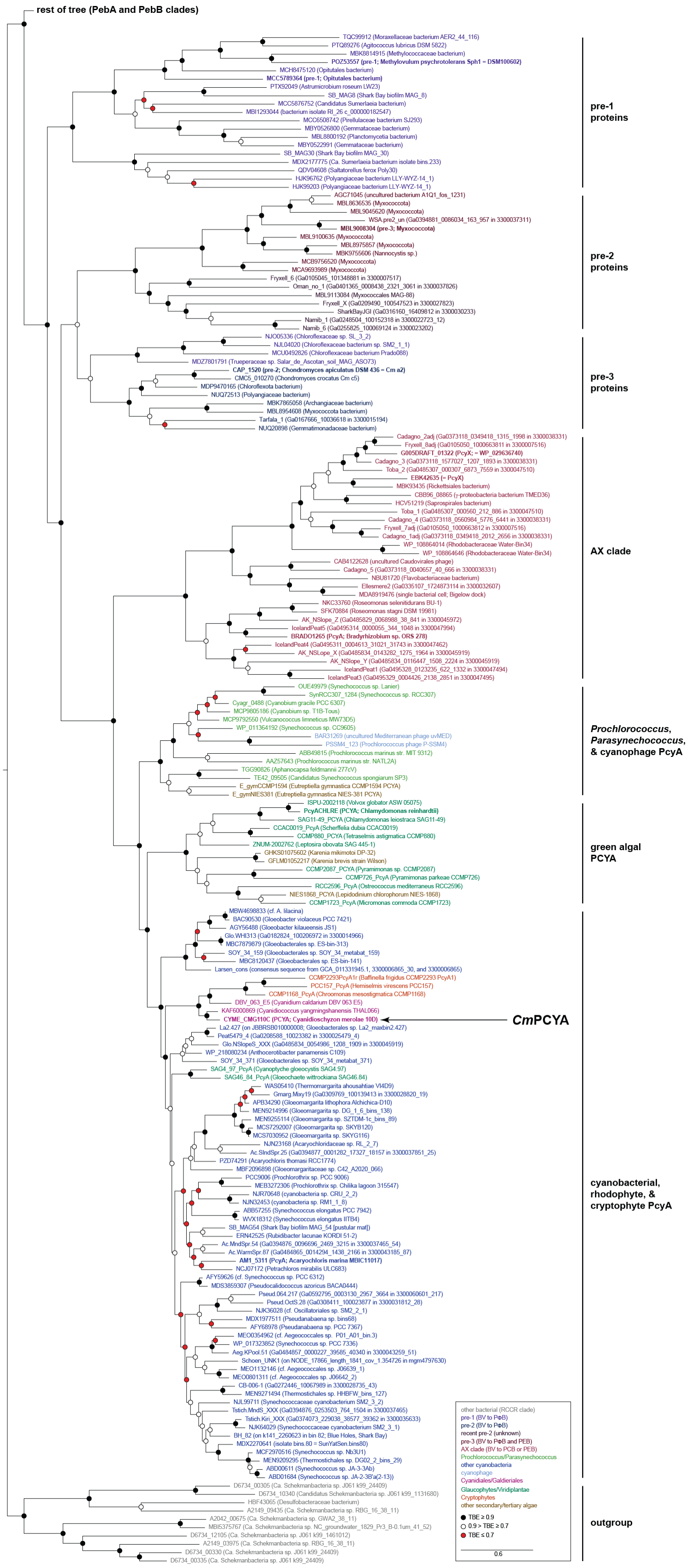


**Supporting Figure S3. Phylogenetic analysis of the PcyA lineage.** The section of the tree containing pre-PcyA, PcyA/PCYA, and PcyX is shown, with the rest of the tree presented in

Supporting Figure S2. Details of tree construction are presented in the Methods. TBE, transfer bootstrap expectation.

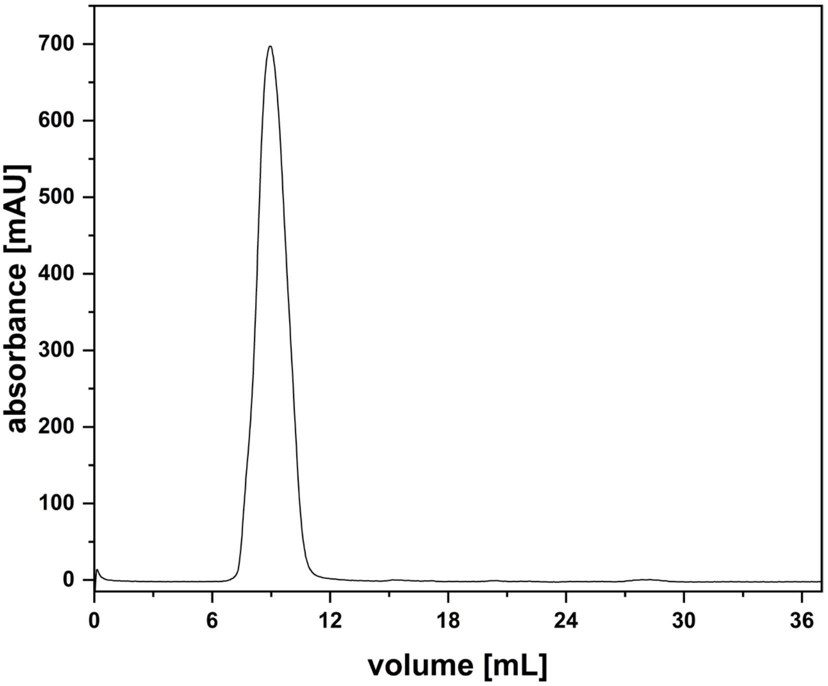


**Supporting Figure S4. Size exclusion chromatography of bovine serum albumin (BSA).**

Size exclusion chromatography was performed using the ÄKTA™ pure 25 purification system equipped with a Superdex™ 75 10/300 GL, equilibrated with “Assay buffer”. Absorbance was continuously measured at 280 nm.

**Supporting Table S1.** ***Galdieria sulphuraria*** **growth media and Trace elements composition.**

| **Component** | **Concentration** |
|---|---|
| $CaCl_2 \times 2\ H_2O$ | 0.14 mM |
| FeNa-EDTA | 0.04 mM |
| $KH_2PO_4 \times 2\ H_2O$ | 2.2 mM |
| $MgSO_4$ | 1.22 mM |
| NaCl | 0.34 mM |
| $(NH_4)_2SO_4 \times 7\ H_2O$ | 11.35 mM |
| Glucose | 25 mM |
| Trace elements | 2 mL/L |

| **Trace elements** | **Concentration** |
|---|---|
| $CoCl_2 \times 6\ H_2O$ | 0.17 mM |
| $CuSO_4 \times 5\ H_2O$ | 0.32 mM |
| $H_3BO_3$ | 46.26 mM |
| $MnCl_2 \times 4\ H_2O$ | 9.2 mM |
| $NaVO_3 \times 4\ H_2O$ | 0.21 mM |
| $(NH_4)_6Mo_7O_{24} \times 4\ H_2O$ | 0.85 mM |
| $ZnSO_4 \times 7\ H_2O$ | 7.7 mM |

**Supporting Table S2. Proteins exclusively identified in the isomerase-active enriched fraction of *Galdieria sulphuraria*, absent in the proteome of *Cyanidioschyzon merolae*.**

| Accession (UniProt) | Annotation | Size (kDa) | Signal peptide? |
|---|---|---|---|
| M2X4W0 | Uncharacterized protein | 13 | No |
| M2XTV5 | BAR domain-containing protein | 34 | No |
| M2XZL8; M2Y956 | Formamidase | 43; 51 | No; No |
| M2VXK0 | Glutathione S-transferase | 25 | No |
| M2X7L6 | Uncharacterized protein | 53 | No |
| M2WZJ6 | BAR domain-containing protein | 34 | No |
| M2W312 | Uncharacterized protein | 27 | Yes |
| M2Y7Y6 | Ribonuclease | 69 | No |
| M2VY74 | Cyclase family protein | 35 | No |
| M2Y9J5 | Glycerol dehydrogenase | 42 | No |
| M2W040 | VHS domain-containing protein | 55 | No |
| M2W7C7 | Acylaminoacyl-peptidase-related protein | 109 | No |
| M2W3V0 | Histone H1 | 26 | No |
| M2WZU6 | Formamidase | 41 | No |
| M2XGU0 | Carboxymethylenebutenolidase | 31 | No |
| M2W3T8 | Oxidoreductase family protein | 40 | No |
| M2X4X4 | Uncharacterized protein | 21 | No |
| M2Y389; M2Y3M5 | LAO/AO transport system kinase isoform 2 | 39; 44 | No; No |
| M2X690 | MARVEL domain-containing protein | 22 | No |
| M2VUV4 | Replication factor A1 | 14 | No |
| M2XZV1 | 2-dehydropantoate 2-reductase | 38 | No |
| M2XXE2 | Alpha-mannosidase | 113 | No |
| M2WYU9 | Uncharacterized protein | 14 | Yes |
| M2XTZ5 | Polysaccharide deacetylase | 36 | No |
| M2XQJ6 | Uncharacterized protein | 29 | No |
| M2XUP9; M2XVD8 | Uncharacterized protein | 33; 35 | Yes; Yes |
| M2XVH6 | Uncharacterized protein | 36 | No |

**Supporting Table S3. Comparison of phycobiliprotein lyases in *Galdieria sulphuraria* and *Cyanidioschyzon merolae*.**

| Lyase | Accession (NCBI) | % identity |
|---|---|---|
| CpcE | GJD06583.1 (*G. sulphuraria*)<br>XP_005534793.1 (*C. merolae*) | 38.28 |
| CpcF | XP_005702676.1 (*G. sulphuraria*)<br>XP_005534794.1 (*C. merolae*) | 35.17 |
| CpcS | GJD07013.1 (*G. sulphuraria*)<br>XP_005538397.1 (*C. merolae*) | 51.43 |
| CpcU | YP_009051127.1 (*G. sulphuraria*) | / |
| CpcT | XP_005709309.1 (*G. sulphuraria*)<br>XP_005536644.1 (*C. merolae*) | 44.66 |
